# PASP — a whole-transcriptome poly(A) tail length determination assay for the Illumina platform

**DOI:** 10.1101/060004

**Authors:** Botond Sipos, Adrian M Stütz, Greg Slodkowicz, Tim Massingham, Jan Korbel, Nick Goldman

## Abstract

The poly(A) tail, co-transcriptionally added to most eukaryotic mRNAs, plays an important role in post-transcriptional regulation through modulating mRNA stability and translational efficiency. The length of the poly(A) tail is dynamic, decreasing or increasing in response to various stimuli through the action of enzymatic complexes, and changes in tail length are exploited in regulatory pathways implicated in various biological processes.

To date, assessment of poly(A) tail length has mostly relied on protocols targeting only a few transcripts. We present PASP (‘<u>p</u>oly(<u>A</u>) tail <u>s</u>equencing <u>p</u>rotocol’), a whole-transcriptome approach to measure tail lengths — including a computational pipeline implementing all necessary analyses. PASP uses direct Illumina sequencing of cDNA fragments obtained through G-tailing of poly(A)-selected mRNA followed by fragmentation and reverse transcription.

Analysis of reads corresponding to spike-in poly(A) tracts of known length indicated that mean tail lengths can be confidently measured, given sufficient coverage. We further explored the utility of our approach by comparing tail lengths estimated from wild type and Δ*ccr4-1/pan2* mutant yeasts. The yeast whole-transcriptome tail length distributions showed high consistency between biological replicates, and the expected upward shift in tail lengths in the mutant samples was detected. This suggests that PASP is suitable for the assessment of global polyadenylation status in yeast.

The correlation of per-transcript mean tail lengths between biological and technical replicates was low (higher between mutant samples). Both, however, reached high values after filtering for transcripts with greater coverage. We also compare our results with those of other methods. We identify a number of improvements that could be used in future PASP experiments and, based on our results, believe that direct sequencing of poly(A) tails can become the method of choice for studying polyadenylation using the Illumina platform

## Introduction

Polyadenylation — the cleavage of the nascent mRNA and the addition of a tract consisting of adenine bases in a non-template-dependent manner — is a post-transcriptional modification characteristic of the majority of eukaryotic transcripts [1–3]. It is performed by a well-characterised protein complex (reviewed in [1]) initiated by a main poly(A) site signal and modulated by various upstream and downstream regulatory elements (reviewed in [4,5]). Both the presence of the poly(A) tail and variation of its length in response to various stimuli are important in numerous biological processes.

There can be several main signal elements within a single transcript defining distinct poly(A) sites (PASs) with different affinity [2,3]. Since the PASs define the actual end of the transcript, differential PAS usage contributes to the processes increasing the complexity of eukaryotic transcriptomes. The differential usage of alternative PASs under different conditions is modulated by various protein factors and chromatin structure [3], and as such it is a part of the repertoire of post-transcriptional regulatory mechanisms of gene expression along with splicing [6]. The length of the poly(A) tail (PAT) itself also plays an important role in post-transcriptional regulation of gene expression and various cellular processes such as mRNA export [7] and stability [5,8,9], translational efficiency [5], cell cycle regulation [10] and microRNA action [11].

The poly(A) tails are bound by poly(A) binding proteins (PABs) during their synthesis in the nucleus. After export to the cytoplasm the PABs associate with the eIF4-G translation initiation factor binding the 5′ 7-methylguanylate cap (m^7^G). This creates a circular structure which protects the mRNA from the cytoplasmic exonucleases, giving rise to the well-established effect of the tail length on mRNA stability [12,13]. The m^7^G and the PAT also synergistically increase translational efficiency [14], further emphasizing the role of the circular structure. While the evidence for the enhancing effect of the PAT on translational efficiency is ample [8,15], the recent high-throughput study by [12] suggests that the role the tail length plays in the regulation of translational efficiency is limited to certain cellular contexts, such as early embryogenesis. For example, [12] found no correlation between tail length and translational efficiency in *Saccharomyces cerevisiae*.

Through its role in the cellular processes above, changes in the length of the PAT are involved in a range of important biological processes such as inflammation [5], embryogenesis [5,12], axonal transport [5], circadian rhythm [16] and DNA damage response [17]. The initial length (~250 bases in humans, ~70-80 in yeast according to [9]) to which the poly(A) polymerase extends the tail is regulated by the affinity and homopolymer structure of the PABs. After synthesis, the tail length is modified by various adenylases and deadenylases, with the enzymes and mechanisms involved showing considerable variability between different organisms [9,18]. In the case of *S. cerevisiae*, the tail is tailored to a message-specific length by the nuclear PAN deadenylase complex, governed by regulatory elements located in the 3′-UTR [18,19]. Throughout the lifetime of the mRNA in the cytoplasm the tail is gradually shortened by the Ccr4-Not complex. Due to this and similar processes, the tail length in total mRNA usually differs from the default length (median tail length 75-100 in human; mean 33 in yeast according to [12]).

The genome-wide study of alternative polyadenylation has been stimulated by the development of assays based on high-throughput sequencing (reviewed in [2]). Regular RNA-seq is inefficient for the study of PASs and PATs, as only a small fraction of the reads overlap with these regions for reasons that are not fully understood [20,21]. Consequently, assays have been developed that compensate for this through enrichment techniques. An additional complication is that on the Illumina sequencing platform reads overlapping PATs have low quality values [21], suggestive of unreliable base calls. Likely causes of this issue are mispriming of the sequencing oligo and increased phasing noise due to polymerase slippage [21]. Both issues are related to the low sequence complexity of the tail; hence some protocols, such as SAPAS [20], introduce random mutations in the sequencing library in order to mitigate this.

The development of high-throughput PAT length assays has been slower than the development of protocols targeting PASs. Until the recent publication of the PAL-seq method by [12], high-throughput poly(A) assays were based on fractionation of mRNAs according to the tail length using affinity chromatography followed by microarray analysis [19,22]. Hence, most of the literature regarding PAT length relates to low-throughput experiments using various techniques targeting specific transcripts [23] and based on reverse transcription and PCR using transcript-specific and tail-specific primers. The tail-specific primers can be simple oligo dTs (as in the RACE-PAT assay [23]) or primers targeting tags at the end of the tail engineered through ligation (LM-PAT [23]), extension by the Klenow fragment (E-PAT [24]), or G-tailing [25]. These methods are well-established; however, they are impractical for large-scale studies.

The high-throughput PAL-seq assay [12] is a customized experiment on the Illumina sequencing platform involving a “T-filling” step similar to [21] using a mixture of dTTP and biotinylated dUTP and measurement of fluorescence of labeled streptavidin bound to the clusters. After normalization based on signal from spike-ins with poly(A) tracts of known length, the measured fluorescence is converted into estimated tail lengths. The PAL-seq assay has good reproducibility (with a correlation of 0.93 between biological replicates), but considering the complexity of the protocol there is still advantage to be gained from alternative high-throughput PAT length assays, preferably based on direct sequencing of the tail.

In this study we explore PASP, an alternative high-throughput Illumina-based PAT assay based on the (low-throughput) G-tailing approach of [25]. In brief, our approach is based on G-tailing poly(A)-selected mRNA followed by fragmentation and reverse transcription using a mixture of primers targeting the poly(A)/G-tag junction and the transcript-end/poly(A) transition. The resulting cDNA fragments are sequenced using a standard Illumina RNA-seq protocol.

We validate PASP by experiments performed on wild type (WT) *S. cerevisiae* and Δ*ccr4-1/pan2* mutants. In addition to yeast being the most popular model organism for studying polyadenylation and developing novel PAT assays, it is known that the Δ*ccr4-1/pan2* mutant has more homogeneous and longer tails which better reflect the default size after synthesis in the nucleus [17,19]. We use the prior expectation this establishes regarding both the direction and magnitude of shift in tail length when compared to the WT to assess the performance of PASP. We also compare PASP to the state-of-the art Illumina-based (PAL-seq) and microarray-based (PASTA) assays.

During the preparation of this manuscript, an approach similar to PASP was published [26]. Due to the time and expense needed to compare this to our protocol, we leave this for a future study.

## Materials and methods

### RNA isolation

WT yeast (BY4741, which is derived from S288c) and Δ*ccr4-1/pan2* [19] were single colony-streaked on YPAD-agar plates, and two individual colonies each were inoculated in 2ml YPAD medium and grown o.n. at 30°C at 180rpm. OD was measured and 7×10^7^ cells were pelleted, resuspended in 2ml Y1 medium (2*μ*l beta-ME, 150*μ*l zymolase) and incubated for 30min at 30°C in a waterbath. After centrifugation, the pellet was lysed in buffer RLT and processed according to the yeast protocol of the Qiagen RNeasy kit. Yield and quality of total RNA was measured with Nanodrop and confirmed by Agilent RNA Nano chip (RIN 10). A DNase digestion was performed on 20*μ*g totalRNA using Turbo DNase (Life Technologies), followed by an RNeasy column cleanup step and elution into 30*μ*l nuclease free water (Ambion).

### Poly(A) selection

Two rounds of poly(A) selection were performed using the TruSeq RNA Sample Preparation kit (Illumina) with several modifications. In detail: 50*μ*l of DNase-digested totalRNA was incubated with 50*μ*l RNA Purification beads (oligo-dT) for 5min at 65°C, cooled to 4°C, and then incubated for 5min at RT in a DNA Engine tetrade 2 (Biorad) thermocycler. Afterwards, beads were separated with a magnet and washed with 200*μ*l Bead Wash buffer, and then eluted in 50*μ*l elution buffer for 2min at 80°C. On RT, 50*μ*l of binding buffer were added, placed on a magnet and washed with 200*μ*l Bead Wash buffer, and again eluted in 50*μ*l elution buffer for 2min at 80°C. The supernatant containing the mRNA was transferred into a 1.5ml Low Binding tube (Eppendorf) and quantified with the Qubit RNA assay.

### Spike-in control

20*μ*l of spike-in *in vitro* transcripts with a known A_42_ stretch [27] were added. A cleanup step was performed using the RNeasy MinElute Cleanup Kit (Qiagen) according to protocol, eluting with 14*μ*l water.

### G-tailing and fragmentation

The USB Poly(A) Tail-Length Assay kit (Affymetrix) was used with several modifications to add a short G stretch to the 3′ end of the mRNA. In detail, 14*μ*l mRNA (with spike-in) were incubated with 4*μ*l of 5x Tail buffer mix and 2*μ*l 10x Tail Enzyme mix at 37°C for 1h and stopped by addition of 2*μ*l 10x Tail stop solution. After an RNeasy Minelute cleanup step and elution into 18*μ*l of water, chemical fragmentation using the NEBnext Magnesium RNA Fragmentation kit (NEB) was performed in 20*μ*l for 3.5min at 94°C. Afterwards, the reaction was immediately put on ice, and 2*μ*l of RNA Fragmentation Stop Solution was added. Another RNeasy Minelute cleanup step was performed, eluting into 15*μ*l of water.

### cDNA synthesis

1*μ*l TTTTVN (16.7*μ*M) and 1*μ*l CCCCCCTT (50*μ*M) custom primers (both Sigma) were added to the 15*μ*l fragmented RNA, primed by incubation at 65°C for 5min and put on ice. First strand cDNA synthesis was performed using the First Strand Master Mix (Illumina TruSeq RNA kit) and SuperscriptII enzyme (Life Technologies) by incubating at 25°C for 10min, 42°C for 50min and 70°C for 15min, and cooled to 4°C in a MJ-Mini thermocycler (Biorad). Second strand synthesis was performed by addition of 25*μ*l Second Strand Master Mix and incubation for 1h at 16°C.

### NGS library preparation

An AMPure XP (Beckman Coulter) cleanup step was performed by adding 90*μ*l AMPure XP beads, vortexing and incubating for 15min at RT. After separation on a magnet, the supernatant was removed and the beads were washed twice with 200*μ*l 80% ethanol. After air drying for 10min at RT, cDNA was eluted in 50*μ*l Resuspension buffer and stored at −20°C. End repair was performed by adding 10*μ*l Resuspension buffer and 40*μ*l Endrepair mix and incubating at 30° C for 30min at 750rpm in a thermomixer (Eppendorf). After another AMPure XP cleanup step using 160*μ*l beads and eluting into 17.5*μ*l Resuspension buffer, the A-overhang addition step was performed. 12.5*μ*l A-tailing mix were added and incubated at 37°C for 30min at 750rpm, followed by 5min at 70°C and put on ice. Adapter ligation used 2.5*μ*l of a 1:40 dilution of Adapter oligo mix (Agilent Sureselect v4 kit), 2.5*μ*l Resuspension solution and 2.5*μ*l DNA ligase mix, incubated at 30°C for 10min in a water bath and stopped by addition of 5*μ*l Stop ligase mix. The adapter-ligated cDNA library was cleaned up by two rounds of AMPure XP using first 42*μ*l of AMPure XP beads and eluting in 50*μ*l Resuspension buffer and then using 50*μ*l AMPure XP beads and eluting in 40*μ*l Resuspension buffer.

Next, a size selection step was performed to create four technical replicates between 300-325bp, 325-350bp, 400-425bp and 425-450bp. For this, the purified adapter ligated cDNA was loaded on a 1.5% Agarose gel and separated for 2h at 120V. Using SybrSafe (Invitrogen) as dye and a Dark Reader transilluminator (Clare Chemical Research), narrow bands were cut using 100bp marker as guidance and gel extracted using the Nucleospin Gel and PCR Cleanup kit (Macherey-Nagel) and eluted in 22*μ*l NE buffer. Two sequential rounds of PCR were performed to introduce barcodes and amplify the cDNA library. First, the Sureselect primer and Sureselect ILM Indexing Pre-Capture PCR reverse primer were used with the Phusion HF 2x mix in 50*μ*l in a MJ Mini thermocycler as follows: 2min 98°C, followed by 5 cycles of 30sec 98°C, 30sec 65°C, 1min at 72°C, and a final 72°C step for 10min. Next, an AMPure XP cleanup step was performed using 90*μ*l AMPure XP beads and eluting into 23*μ*l water. Afterwards, a second PCR was performed using the Sureselect ILM Indexing Post Capture Forward PCR primer and a different PCR Primer Index primer/sample together with the Phusion HF 2x mix in 50*μ*l in a MJ Mini thermocycler as follows: 2min 98°C, followed by 13-15 cycles of 30sec 98°C, 30sec 57°C, 1min at 72°C, and a final 72° C step for 10min. After a final AMPure XP cleanup step using 90*μ*l beads, the final library was eluted in 20*μ*l water, quantified with the Qubit HS dsDNA kit and pooled equimolarly.

### Illumina sequencing

All 16 pooled barcoded libraries were sequenced on one HiSeq2000 lane in 2 × 101bp mode. Raw data have been deposited to Array Express under accession number E-MTAB-2456.

### Data analysis

The data analysis pipeline with documentation, logs and selected raw results (see also **Supporting Text S1**) is available at https://github.com/bsipos/paper-pasp. Individual primary data analysis steps implemented in specialised tools written in Python (under directory `patsy/`) and downstream data analysis steps implemented in R scripts (under `scripts/`) can be invoked through documented make targets.

The first primary data analysis step is the classification of read pairs into “G-tail” pairs, based on the presence of a sequence motif characteristic to the poly(A) tail/oligo(G)-tag junction in the first 14aligned bases of one of the reads, or “NVTR” read pairs otherwise. For G-tail reads we require a minimum G-tag length of 3 and we allow for a maximum of 6 Ns in the first 14 bases. The G-tail pairs represent those that will, ideally, contain the complete PAT in one read and mappable sequence in the paired read that will unambiguously identify the transcript. We map the G-tail read pairs to the SGD *S. cerevisiae* reference transcriptome [28] using Bowtie2 [29] with parameters --`ignore-quals` --`sensitive -I 0-X 500 --no-contain`, chosen to perform a stringent alignment of the read anchoring the fragment to the respective transcript.

All non-G-tail read pairs are classified as NVTR reads, representing those targeting the transcript-end/poly(A) junction as well as “noise” reads most likely originating from fragments that are products of off-target priming. The NVTR reads are used to assess the insert size distributions of the technical replicates and other quality control measures. They are mapped to the SGD transcriptome using Bowtie2 with parameters `--ignore-quals --sensitive-local-I 0-X 500 --no-mixed --no-discordant --no-contain`, aiming for a less stringent alignment of both reads corresponding to the fragment. The classification and mapping analysis steps are implemented in the `patsy-align` tool.

The resulting G-tail and NVTR alignments are parsed into G-tail and NVTR fragments, respectively, using the `patsy-parse` tool. For each transcript, we tabulate the distribution of the number of A bases until the first non-A base after the G-tag in the G-tail reads overlapping the PAT (the “tail run” distribution). Insert size distributions are assessed based on the mapped NVTR fragments.

Using the `patsy-spike` tool we extract the reads mapped to the spike-in standards, identify the reads overlapping the poly(A) tracts with known lengths and plot the distributions of number of A bases until the first 1-5 non-A bases. Comparison of the results obtained for this range of thresholds, necessary because of the low quality scores of many reads, led us to choose a criterion of 1 non-A base for defining the PAT length.

The last step of the primary data analysis is to plot the per-transcript and global tail run distributions from wild type (WT1 and WT2) and mutant (MUT1 and MUT2) samples and to test the significance of differences using Mann-Whitney U-test (with a minimum sample size of 30).

Downstream analysis steps (invoked through other analysis pipeline make targets) perform filtering by G-tail fragment coverage, plotting of the correlation between biological replicates and technical replicates of the size selection step (above) as a function of different coverage thresholds, plotting and clustering of the per-transcript tail run distributions and correlation of thresholded per-transcript mean tail run lengths with results generated by the PAL-seq [12] and PASTA [19] assays (see the pipeline documentation).

## Results and discussion

### Characteristics of aligned fragments

The number of NVTR fragments was consistently 2.6-7.4 times higher than the number of G-tail fragments in all technical replicates. Also, the majority of the reads overlapping the poly(A) tails were sequenced as second reads, indicating that there were issues with detecting clusters having the repetitive tails as first reads (see alignment reports listed in **Supporting Text S1**).

The NVTR fragments also had a higher mapping rate (49-66%) than the G-tail fragments (26-39%). NVTR fragments mapped to various locations within the transcripts (see the report files listed in **Supporting Text S1**), and for that reason we did not further investigate whether these fragments reliably define individual polyadenylation sites.

### Quantification of tail run “slippage” using spike-ins

The poor quality of reads overlapping the poly(A) tails and the “drift” of the sequenced tail length is a known issue [21] which can be clearly observed in the reads mapping to the poly(A) tracts in our spike-ins (see the file linked from **Supporting Text S1**). However, the mean tail run length from these 53558 reads is close to the true length of the poly(A) tracts (44.56 ± 16.11 cf. 42; see **Figure 1**). This suggests that given enough coverage of G-tail reads there is sufficient signal for characterising the tail length, as with increasing sample size the mean of the tail run distribution (including slippage noise) gets closer to the true tail length. The variance of the slippage noise distribution, however, also implies that filtering out transcripts with low G-tail fragment coverage is an essential step in quality control of PASP data.

**Figure 1.**
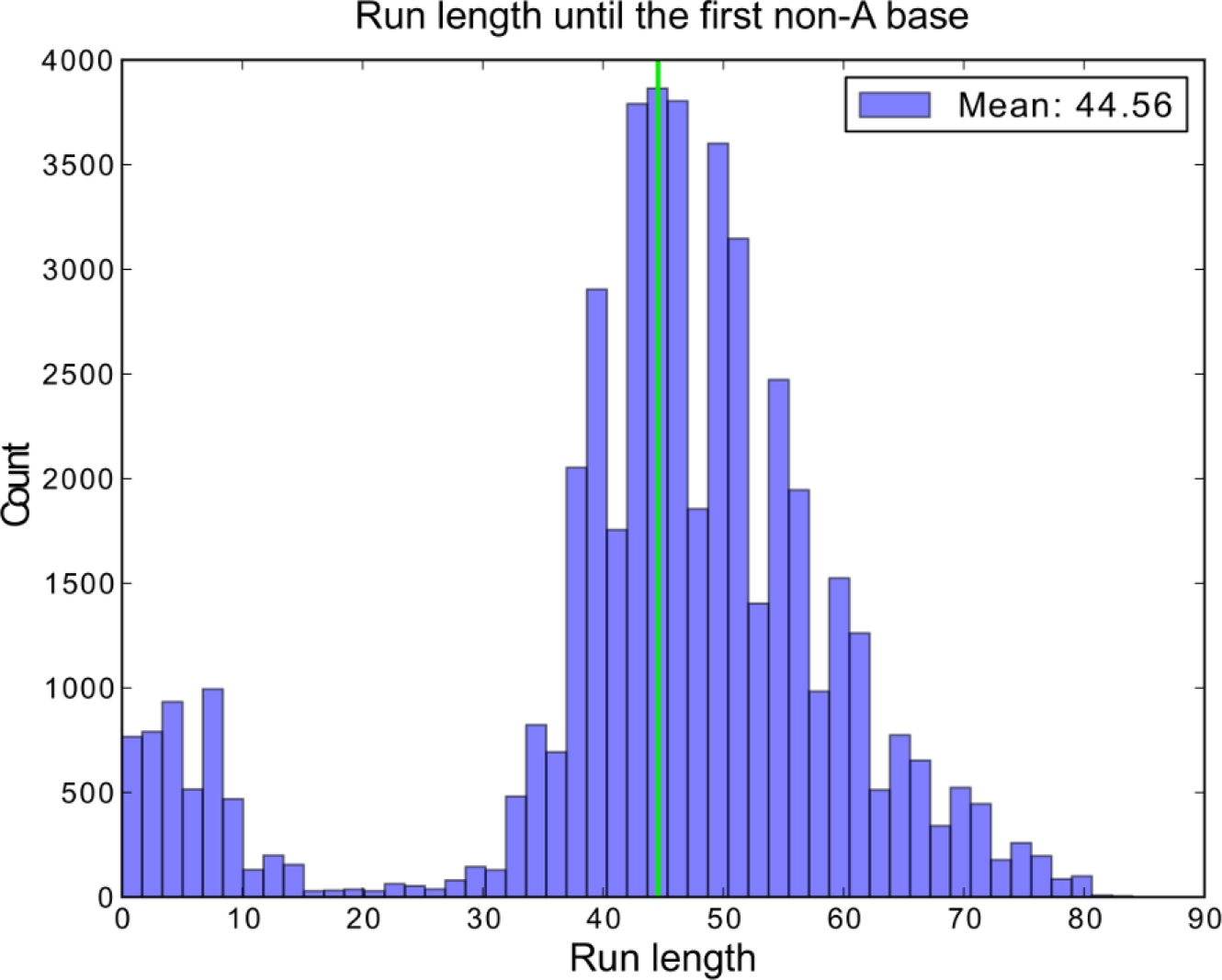
The distribution of the number of aligned A bases until the first non-A base (“tail runs”) in the reads mapping to the poly(A) tracts of known length of the spike-in standards. The vertical green line marks the mean run length from all 53558 reads (44.6); the true length is 42.

### Global shift in polyadenylation status in Δ*ccr4-1/pan2* mutants

The distribution of tail run lengths tabulated from the whole WT and mutant transcriptomes (**Figure 2A**) showed high consistency between biological replicates (compare **Figure 2** with **Supporting Figure S1**). Both distributions are unimodal with a fat tail to the right. The right tail of the distribution is truncated in the case of the mutant samples as the largest measurable tail run is limited by the read length; the much lesser degree of truncation for the WT samples suggests that the read length of 101bp is capturing the full length of virtually all their poly(A) tails.

**Figure 2.**
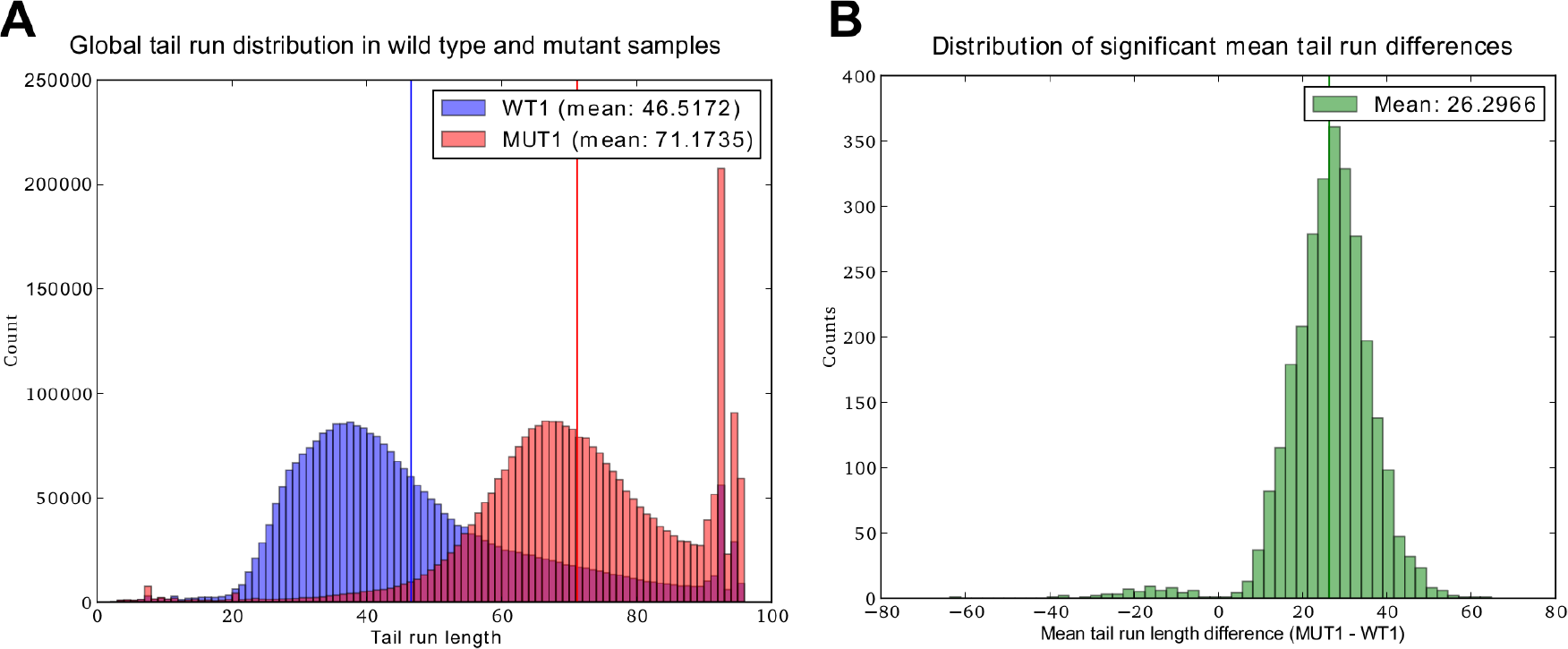
Shift in global polyadenylation status in Δ*ccr4-1/pan2* mutants. **A**. Distribution of tail run lengths from all transcripts in the wild type (WT1, blue) and the mutant (MUT1, red) samples. The coloured vertical lines indicate the means of the respective distributions. **B**. The distribution of significant (Mann-Whitney U-test, *p* < 0.05) per-transcript mean tail run length differences (MUT1 - WT1).

We consistently observe an upward shift in the tail length in the Δ*ccr4-1/pan2* mutants: in WT samples the global mean of tail run lengths is consistently around 46, which is approximately in line with values reported by other studies (median tail length of 33 reported by [12]), while in the mutants the mean length is around 71. Moreover, the global shift in tail run length (+25) is consistent with the mean of per-transcript tail run shifts (compare **Figure 2B** with **Figure 2A**). These observations are in accord with expectations, given that the mutant is supposed to have longer tails (>55 according to [19]) due to the lack of nuclear and cytoplasmic deadenylase activity. This suggests that PASP is suitable for the assessment of global polyadenylation status in yeast.

### Coverage-dependent consistency of per-transcript tail run distributions

Despite the consistent global tail run distributions, the correlation of per-transcript mean tail run lengths between biological replicates is not high, both in the case of wild type and mutant samples (WT1 vs. WT2: *r*=0.25; MUT1 vs. MUT2: *r*=0.46). However, the correlation increases after applying coverage filtering with increasing thresholds. After filtering for minimum G-tail coverage of 400 fragments, we are left with 1211 transcripts (out of 3951 detected overall) in wild type and 673 (out of 4646 detected) in the mutant samples, with per-transcript mean tail run correlations of 0.54 and 0.95, respectively. At a coverage threshold of 1000 the correlation reaches high values both between wild type and mutant samples (WT1 vs. WT2: r=0.73; MUT1 vs. MUT2: r=0.98; **Figure 3A**), but we lose a greater proportion of the detected transcripts (465 transcripts retained in the WT samples; 315 transcripts retained in the mutant). This behavior is consistent with the observations drawn above from the analysis of spike-in reads, and also explains the consistency of the global tail run distributions which are necessarily dominated by more abundantly expressed transcripts.

**Figure 3.**
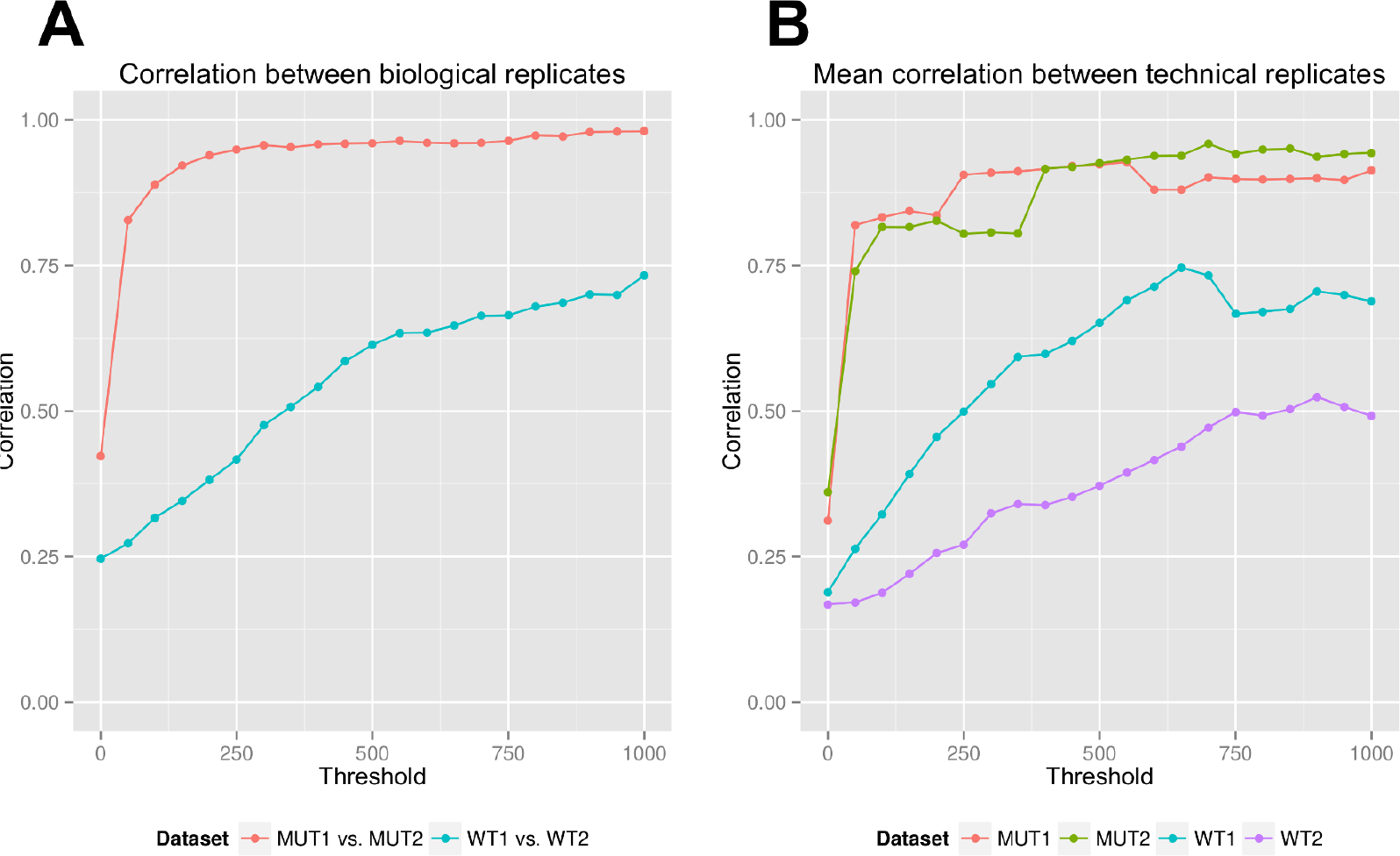
Pearson correlation between biological and technical replicates at different G-tail fragment coverage thresholds. **A**. Correlation between biological replicates. **B**. Average correlation between technical replicates from a given sample.

The increase of correlation between mutant and WT biological replicates showed some interesting differences: correlation between the mutant replicates is already high at low coverage thresholds and initially increases with a higher rate; this is, however, coupled with a higher number of discarded transcripts. The higher initial and post-filtering correlation between the mutant replicates likely has both technical and biological causes. First, the tails in mutants are likely to stay closer to their initial length and so are expected to be more homogeneous [17,19]. Second, tails in the mutants are closer to the read length; hence the drift towards longer lengths is expected to have a smaller confounding effect.

The mean correlation between pairs of technical replicates showed a similar behaviour at increasing coverage thresholds in the case of the two mutant samples (**Figure 3B**), both curves being similar in shape to the change in correlation between mutant biological replicates. This suggests that the mutant samples suffer from comparable amount of technical noise and that this is the likely cause of reduced correlations at lower coverage thresholds.

The mean correlation between technical replicates is substantially lower in the case of WT samples (**Figure 3B**), which can be explained by either higher intra-transcript variability or higher levels of technical noise. The correlations are especially low in the the case of WT2. This makes it likely that WT2 suffers from the highest levels of technical noise, which can be partly responsible for the lower correlation between WT biological replicates.

Regardless of low correlation at low G-tail fragment coverage, tail run distributions showed excellent consistency between WT biological replicates in the case of highly expressed transcripts (see examples in **Figure 4**). Most transcripts with high G-tail fragment coverage have longer mean tail runs in the mutant samples (**Figure 4A,B**), with the magnitude of difference being similar to the difference in global mean tail lengths, However, a few transcripts — most notably transcript YER177W (BMH1) — had a short mean tail length in both the WT and mutant samples (**Figure 4C**). This suggests that poly(A) tail length in this transcript might be under another specific regulatory mechanism, potentially worthy of further investigation.

**Figure 4.**
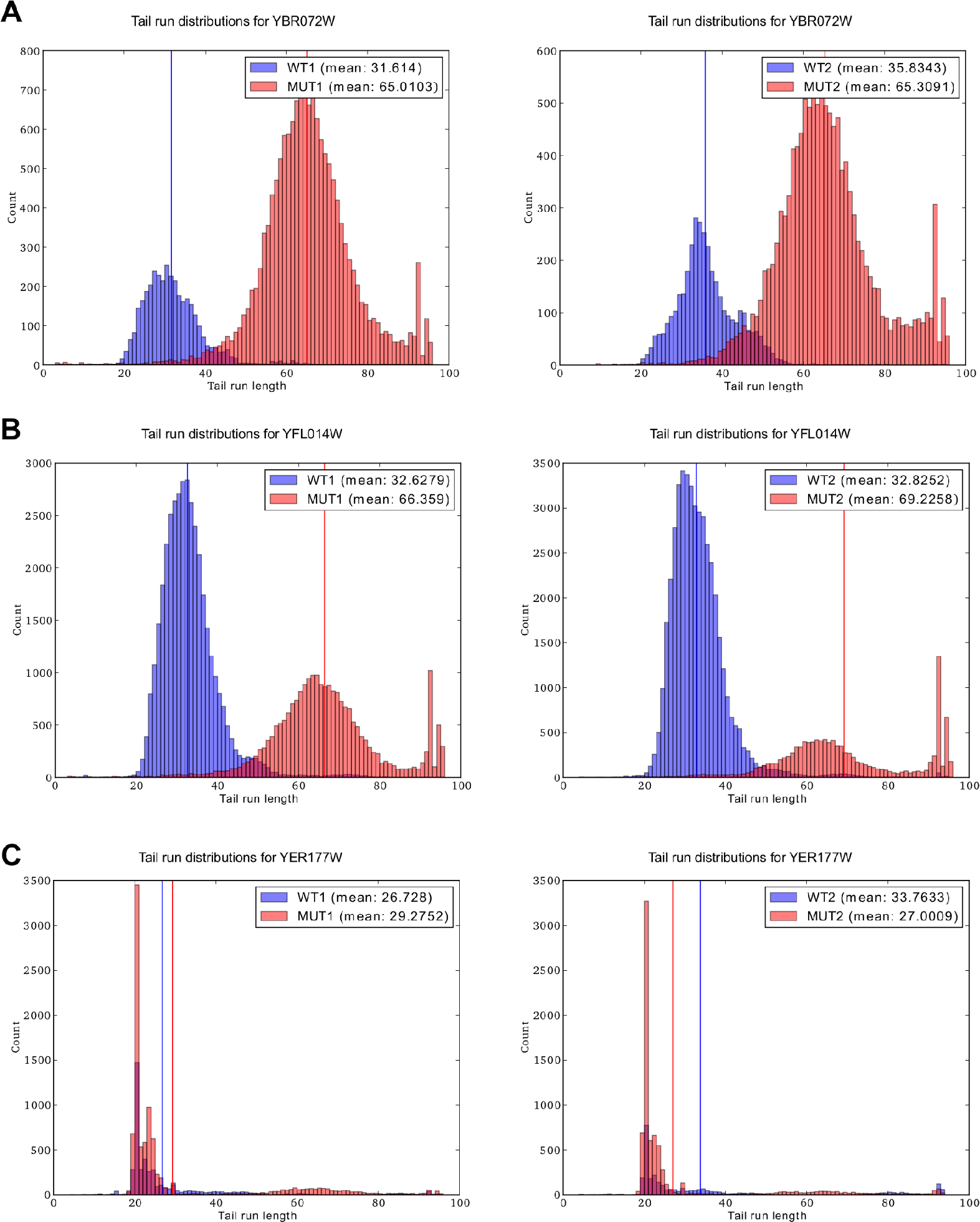
Tail run distributions for selected transcripts with high G-tail fragment coverage in WT1 and MUT1 (left) and WT2 and MUT2 (right) samples. **A**. Small heat shock protein HSP26/YBR072W. **B**. HSP12/YFL014W. **C**. BMH1/YER177W.

The consistency of the tail run distributions for transcripts such as those shown in **Figure 4** suggests that, given sufficient coverage, PASP has the potential to provide a reliable assay of per-transcript mean poly(A) tail length and potentially provide information about the details of the tail length distribution (e.g. multimodality). However, considering the lower number of high-coverage transcripts and the poor correlation with the PAL-seq assay (see below) we refrain from further downstream analyses of the per-transcript tail run lengths in this experiment.

### Correlation between PASP and previous high-throughput assays

We have examined the correlation of WT mean per-transcript tail run lengths for the 465 transcripts with at least 1000 G-tail fragment coverage with results obtained from *S. cerevisiae* by the PASTA [19] and PAL-seq [12] protocols. There was no correlation between the PASP and PAL-seq results (WT1 vs. PAL-seq: *r*=0.07; WT2 vs. PAL-seq: *r*=0.03; **Supporting Figure S2**), despite the high correlation (*r*=0.73) of PASP between biological replicates for the high coverage subset and the high correlation between PAL-seq biological replicates. The correlation between PAL-seq and PASTA results was also weak (r=0.14 for 3631 transcripts; **Supporting Figure S3**), while correlation between PASP and PASTA was negative (**Supporting Figure S4**).

The observed poor correlation between the different assays suggests that the polyadenylation status of the transcriptome might be especially sensitive to growth conditions, or alternatively can be explained by strong and reproducible biases affecting one or more of the assays. Additional experiments will be necessary to further compare these alternative methods.

### The potential of PASP and future directions in studying polyadenylation

We believe that in the future, as sequencing technologies permit longer and more accurate reads, assays based on direct sequencing of the PAT will be the dominant techniques for studying the biology of polyadenylation. We also believe that PASP has the potential to become the assay of choice on the Illumina platform. The results of our pilot experiment suggest several ways to improve its properties. First, we have demonstrated that the reliability of the tail length estimates can be increased simply by increasing sequencing depth, suggesting that by using one Illumina lane per sample it might be possible to reliably assess the polyadenylation status of the complete yeast transcriptome.

Second, adding random barcodes to the sequencing adapter could solve the cluster detection issues and recover many lost G-tail fragments. It may also be possible to ameliorate sequencing problems caused by the repetitive tail by introducing random mutations in the library as has been suggested in the SAPAS method [20] or the NG-SAM approach to sequence assembly [30].

Third, improvements in Illumina chemistries and base calling algorithms might improve the quality of reads overlapping the tail even without any alterations to our experimental protocol. With one or more of these improvements, the PASP approach could provide a practical alternative to more complicated high throughput assays such as PAL-seq or protocols based on the still relatively expensive third-generation sequencing platforms [31].

A shortcoming of PASP is that the maximum tail length which can be detected is limited by the read length. This means that for organisms with potentially longer tails (such as mammals), longer reads are needed in order to achieve sufficient dynamic range. The MiSeq Illumina platform is already capable of generating longer reads suitable for this purpose.

One of the most puzzling results of our pilot study is the low correlation between the PASP and PAL-seq assays, even for the set of high-coverage transcripts and despite excellent reproducibility between biological replicates for both methods. While additional experiments are needed to uncover the source of this discrepancy, this observation suggests that the variability of the polyadenylation status of the yeast transcriptome in response to growth and environmental factors is yet to be uncovered.

## Conclusions

In this study we have presented PASP, a new high-throughput approach to measure poly(A) tail lengths using Illumina short read sequencing. We demonstrated the utility of this approach by estimating and comparing tail lengths from wild type *S. cerevisiae* and a deadenylase-mutant with expected longer tails. We found that, given high-enough coverage, the tail lengths can be confidently measured even in the presence of “slippage” noise.

The distributions of tail run lengths tabulated from the whole wild type and mutant transcriptomes showed excellent consistency between biological replicates, suggesting that PASP is suitable for the assessment of global polyadenylation status in yeast. Correlation of per-transcript mean tail lengths between biological and technical replicates was high only for transcripts with high G-tail coverage, implying that filtering-out transcripts with low coverage is an essential step in quality control of PASP data.

Based on our results we believe that direct sequencing of poly(A) tails will be the method of choice for studying polyadenylation and that after implementing the suggested improvements PASP has the potential to become an assay of choice for the Illumina platform.

## Acknowledgements

We would like to thank Prof. Thomas Preiss and Stuart Archer for providing the Δ*ccr4-1/pan2* mutant yeast strain, Vicente Pelechano for providing the spike-in standards and helpful discussion in the planning stage and Benjamin Raeder for technical expertise.

BS was funded by a European Molecular Biology Laboratory (EMBL) Interdisciplinary Postdoc (EIPOD) Fellowship under Marie Curie Actions (COFUND). AMS, GS, TM, JK and NG were supported by EMBL. GS is a member of Wolfson College, University of Cambridge. BS and TM)s contributions to the work were concluded prior to their employment at Oxford Nanopore Technologies.

